# Study on the preparation of bovine milk exosomes and the stability of lyophilized powder

**DOI:** 10.1101/2023.10.16.562624

**Authors:** Lu Lu, Chunle Han, Miao Wang, Huanqing Du, Ning Chen, Mengya Gao, Na Wang, Dongli Qi, Wei Bai, Jianxin Yin, Fengwei Dong, Tianshi Li, Xiao Hu Ge

## Abstract

Bovune milk exosomes (MK-exo) have bioactive effects on the health of consumers and can also serve as excellent drug delivery vehicles. However, long-term preservation and transportation remain major challenges at present. In this report, high-quality MK-exo are prepared on a large scale. We then developed a lyophilization method belonging to MK-exo, and performed acceleration and long-term stability experiments on the lyophilized MK-exo to evaluate the effect of lyophilization. The results showed that the lyophilized MK-exo can be stored at room temperature for at least 3 months and 2-8°C℃ for 15 months. These findings provide data support for further understanding of the properties and applications of MK-exo.

## 1. INTRODUCTION

Exosomes are a subset of extracellular vesicles (EVs) ranging in size from 40-150 nm(Kalluri 2016, Li et al. 2019). Exosomes isolated from cow’s milk are MK-exo (Munagala et al. 2016) which carry some active substances themselves (Chen et al. 2010, Hata et al. 2010, Zeng et al. 2019). And they have application potential in the food, cosmetics and pharmaceutical industries. First, MK-exo can penetrate the intestinal barrier(Hata et al. 2010, Wolf et al. 2015, Rani et al. 2017) and have important functions such as immune regulation, anti-bacterial infection, anti-oxidative(Zhong et al. 2023), which suggests that they can be used in the development of functional foods and play a beneficial role in human health at multiple levels, including intestinal health, bone/muscle metabolism, and microbiota regulation(Suharta et al. 2021). In addition, MK-exo are beneficial to intercellular adhesion and can penetrate mouse skin, indicating that MK-exo can penetrate the skin barrier and have the effect of repairing it(B. et al. 2022), which shows its potential for use in cutaneous aesthetics(Xiong et al. 2021). Meanwhile, MK-exo successfully loaded with mRNA(Zhang et al. 2023), paclitaxel (Agrawal et al. 2017) and doxorubicin (Pullan et al. 2022), demonstrating their excellent drug delivery capabilities and application potential in the pharmaceutical industry (Del Pozo-Acebo et al. 2021).MK-exo are derived from bovine milk that is safe to eat, suggesting that they can be used non-toxically in other foods, cosmetics, and oral medications in the right dose. Several reports have tested MK-exo for toxicity(Munagala et al. 2016, Agrawal et al. 2017, Somiya et al. 2018), and the results have shown its safety. And the results of the safety evaluation test of MK-exo we have done also show that MK-exo are safe(Lu et al. 2023).

Usually, exosomes are obtained from biofluids or supernatant of cell culture media. Among the various sources, bovine milk has the advantage of being cheaper and easier to scale(Haug et al. 2007). Bovine milk production worldwide is very high and generally inexpensive, and the volume of it produced worldwide had reached to around 544 million metric tons by 2022(Statista). Recently the lower price of milk in New Zealand was about $0.4067 per litre, and the higher price in Japan was $0.8138 per litre(CLAL). At the same time, the price per liter of cell culture medium ranges from a few hundred to thousands of dollars(Scientific, Technologies). In simple terms, purifying exosomes from bovine milk saves the time and cost of cell culture, and does not worry about the risk of cell contamination.

The above findings from the perspectives of functionality, safety and industrialization indicate that MK-exo have industrial value. but there are still some challenges, such as large-scale production, preservation and transportation problems. Size exclusion chromatography (SEC) and tangential flow filtration (TFF) are mature(Blans et al. 2017, Guo et al. 2021, Han et al. 2021, Kim et al. 2021, Gao et al. 2022), and we has been solved large-scale production of exosomes by them. Therefore, preservation and transportation are challenges that become urgent to be solved at present.

A conventional and efficient way to store exosomes is frozen at −80 °C(Zhou et al. 2006, Witwer et al. 2013, Jeyaram et al. 2017, Welch et al. 2017, Thery et al. 2018). Agrawal et al. found that bovine milk-derived EVs can be stored at −80°C for four weeks without any changes in physical properties(Agrawal et al. 2017). Munagala et al. reported no significant changes in the physical and biological properties of MK-exo after 18 months of storage at −80 °C(Munagala et al. 2016). Study by Lorincz et al suggested that storage at −80℃ for 28 days had no significant effect either on neutrophilic granulocytes EV number or size(Lorincz et al. 2014). These data suggest that it is feasible to store MK-exo at −80 °C. But the storage of this mode is limited during transportation, and the use of dry ice causes inconvenience and high cost of transportation(Roy et al. 2004, Gil et al. 2014). Therefore, a simple, convenient and low-cost storage method needs to be urgently obtained.

Lyophilization is a technique that has been used to preserve various types of biological materials, facilitating their storage and transportation without the laborious and expensive cold chain. Different lyophilization methods have been used to improve the long-term stability of different exosomes. Geeurickx et al. reported that the morphology, fluorescence intensity, number, size distribution and density of recombinant HEK293T cell exosomes had no effect after lyophilization(Geeurickx et al. 2019). Le Saux et al. reported that the structure of mMSC-derived EVs was not impacted after freeze-drying(Le Saux et al. 2020). Charoenviriyakul et al. found that B16BL6 cells derived exosomes could be stored stably at room temperature for 4 weeks after lyophilizing, and the proteins and RNA in exosomes were not affected(Charoenviriyakul et al. 2018). In addition, Trenkenschuh et al. reported that mammalian RO EVs after lyophilization was stored at 2-8°C for 1 month, maintaining the original particle size and concentration without cargo loss(Trenkenschuh et al. 2022). Currently, research on lyophilization of exosomes is neither systematic nor in-depth, and data on the stability of lyophilized MK-exo are rarely reported.

In this study, we first found a significant decrease in purity of MK-exo when stored at −80°C for 8 months. Next, we found that lyophilization methods for exosomes of different sources is not universal. We then developed a lyophilization method for MK-exo. In addition, stability experiments verified the stability of lyophilized MK-exo. Here we provide a way for long-term stable storage of MK-exo, solve the problem of their transportation difficulties, and provide convenience for them in various applications.

## 2. MATERIAL AND METHODPreparation of MK-exo by Size exclusion chromatography (SEC) method

The pH of milk was adjusted to pH 4.6 by hydrochloric acid (Merck) followed by centrifugation at 4000×rpm (Beckman) at 4℃ for 30 min. The supernatant was collected and then purified by deep filtration, TFF, and SEC. The isolated MK-exo was sterile filtered through a 0.22 μm filter and stored at −80 ℃ until further use.

### 2.2. Preparation of MK-exo by Density-gradient ultracentrifugation (DC) method

The casein-free whey was purified by deep filtration. Then, Sucrose density gradient centrifugation was performed as described previously(Hata et al. 2010). The isolated MK-Exo was sterile filtered through a 0.22 μm filter and stored at −80 ◦C until further use.

### 2.3. Transmission electron microscope

The MK-Exo (100 μg/mL) was fixed in 2% (w/v) paraformaldehyde at room temperature for 15 min. The mixture (10 μL) was mounted on a formvar-carbon coated grid (Beijing XXBR Technology, Beijing, China) at room temperature for 3 min and then stained by uranyl oxalate solution (4% uranyl acetate, 0.0075 M oxalic acid, pH 7) for 1 min. After the samples were observed by TEM (Hitachi High-Technologies Corporation, Tokyo, Japan).

### 2.4. Analysis of size distribution and particle number

The MK-Exo size distribution and number were determined by NanoFCM (NanoFCM Inc., Xiamen, China) according to the user manual.

### 2.5. Purity analysis

The sample was analyzed using an SEC-1000 column (7.8*150 mm, 7 um, Thermo). The column was eluted at a 0.3 mL/min flow rate with 150 mM NaCl and 20 mM phosphate buffer pH 7.2.

### 2.6. Zeta potential Measurements

The Zeta potential of MK-Exo was measured thrice at 25 °C under the following settings: sensitivity of 85, a shutter value of 70, and a frame rate of 30 frames per second, while ZetaView software was used to collect and analyze the data (Midekessa et al. 2020).

### 2.7. Quantitative assessment of protein concentration

The protein concentration of MK-Exo was measured by BCA Kit (Thermo, Waltham Mass, USA) according to the manufacturer’s instructions.

### 2.8. Proteomic analysis

The LC-MS/MS using Easy NLC 1200-Q Exactive Orbitrap mass spectrometers (Thermo). The nano-HPLC system was equipped with an Acclaim PepMap nano-trap column (C18, 100 Å, 75 μm × 2 cm) and an Acclaim Pepmap RSLC analytical column (C18, 100 Å, 75 μm × 25 cm). One μL of the peptide mix was typically loaded onto the enrichment (trap) column. All spectra were collected in positive mode using full-scan MS spectra scanning in the FT mode from m/z 300-1650 at resolutions of 70000. For MSMS, the 15 most intense ions with charge states ≥2 were isolated with an isolation window of 1.6 m/z and fragmented by HCD with a normalized collision energy of 28. A dynamic exclusion of 30 seconds was applied.

The raw files were searched using Proteome Discover (version 2.4, Thermo) with Sequest as the search engine. Fragment and peptide mass tolerances were set at 20 mDa and 10 ppm, respectively, allowing a maximum of 2 missed cleavage sites. The false discovery rates of proteins and peptides were 1%. DAVID Bioinformatics Resource 2021 (http://david.abcc.ncifcrf.gov/) analyzed the differential expression proteins with recommended analytical parameters to identify the most significantly enriched signal transduction pathways in the data set.

### 2.9. The content of Impurity proteins determination

The whole milk protein and β-lactoglobulin of MK-exo were measured by the RIDASCREEN®FAST Milk kit (R-Biopharm AG, Darmstadt, Germany) according to the manufacturer’s instructions. The casein was measured by the PriboFast^®^ Casein (Bovine milk) ELISA Kit (Pribolab, Qingdao, China). The IgM and IgG were measured by the Strongyloides IgG/IgM ELISA kit (Abcam, Cambridge, UK).

### 2.10. Freeze-drying

MK-exo was lyophilized using methods of lyophilizing exosomes reported by others(L. et al. 2018, Geeurickx et al. 2019).

Then there is the lyophilization method we developed. Trehalose and mannitol were chosen as a cryoprotectant and dissolved into purified MK-exo to obtain a final concentration of 5% (w/v). Typically 1 ml liquid was transferred in 2ml cryogenic vials and frozen in a freeze dryer (SCIENTZ, Ningbo, China).

### 2.11. Water content

The water content of the MK-exo lyophilized powder was repeatedly measured at least twice using a KEM MKA-610 Karl-Fischer moisture titration (Kyoto Electronics, Kyoto, Japan) (Clua-Palau et al. 2020).

### 2.12. Cell line

Caco-2 cells were purchased from the National Institute of Cell Resource, Beijing, China. Caco-2 cells were grown in DMEM (Gibco, CA, USA) with 100 ug/ml streptomycin and 10% FBS). Cells were maintained in a 5% CO_2_ humidified atmosphere at 37°C.

### 2.13. Western blotting

Western blot analyses were performed according to the previously described protocols (Lu et al. 2013). The primary antibodies used in this study were CLD and GAPDH (Abcam). Depending on the primary antibody, the secondary antibodies were either goat anti-rabbit (Abbexa) or goat anti-mouse antibodies (Protientech). The immunoreactive protein was used with ECL (Thermo) to image immunoblots.

### 2.14. Preparation of and 293 cell and MSC exosomes

Expi293F cells were cultured as described previously. The cell suspension was centrifuged at 3,000 g for 10 min at 4°C and the supernatant is harvested.Human mesenchymal stem cell supernatant were prepared as described previously(Wang et al. 2021). The hMSC-exo and 293-exo were purified using the method of density gradient centrifugation from the collected supernatant.

### 2.15. Statistical analyses

The statistical analysis was performed by using GraphPad Prism software (version 8.0, GraphPad Software, San Diego, CA, USA). Data were shown as mean ± standard deviation (SD). The Origin Software was used for the protein quantity accumulation analysis and correlation analysis. Differences were evaluated using Student’s t-test and were considered statistically significant at p < 0.05. The Venny platform (http://bioinfogp.cnb.csic.es/tools/venny/index.html) was used to make Venn diagrams. The bioinfarmatics platform (https://www.bioinformatics.com.cn/) was used to create heat maps.

## 3. RESULT

### 3.1. Preparation and characterization of Milk exosomes

To obtain GMP-compliant MK-exo, we purified milk using the SEC method. Three batches of MK-exo were prepared Transmission electron microscopy results showed a concave tea tray shape (Figure 1A), which is the classic morphology of exosomes. And more than 98% of the particle size is in the range of 40-150nm (Figure 1B). Meanwhile, the proteome results showed positive marker proteins (CD9, CD63, CD81, TSG101, Alix) and no negative marker proteins (Calnexin, Grp94, Cytochrome, GM130) (Figure 1C). The above results showed that MK-exo was indeed prepared by the SEC method.

**Fig 1.**
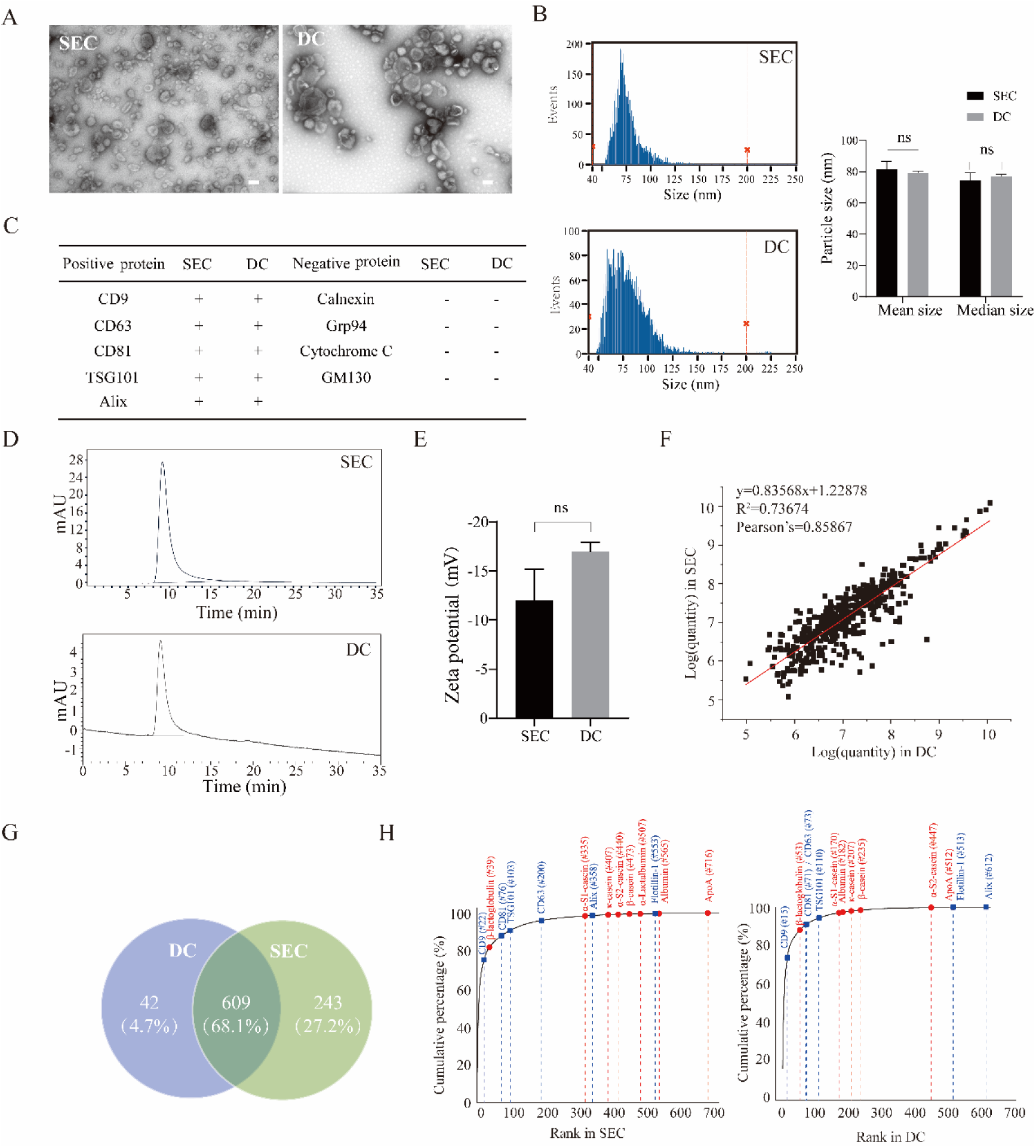
Characterization of MK-exo prepared by SEC and DC. (A-B) The main morphological characteristics and particle size analysis of MK-exo prepared by the method of SEC and DC, respectively; Scale bar = 100 nm; (C) Positive and negative marker proteins of MK-exo; (F) Venn diagram for MK-exo proteome;(G) Protein quantity accumulation analysis between specific surface proteins and major milk proteins (impurity protein) of MK-exo;(H) Protein correlation analysis between the MK-exo prepared by the SEC and DC methods; (D-E)The purity analysis and zeta potential of MK-exo. The error bars represented the standard deviation of three repetitive experiments. ns: There was no significant difference.

DC is considered to be the “gold standard” method for exosome isolation (Gao et al. 2022), so the DC method was used to purify MK-exo to evaluate the quality of MK-exo collected by the SEC method. Next, several metrics continue to be used to evaluate the quality of MK-exo purified by the SEC method. MK-exo prepared by the two methods did not differ significantly in morphology, particle size, and protein marker results (Figure 1A-C). Purity analysis by size-exclusion chromatography both showed only one peak (Figure 1D) and no significant difference in Zeta potential results (Figure 1E). The above results showed that MK-exo isolated by two methods have the same physical properties. Furthermore, the Venn diagram shows that the MK-exo prepared by the SEC method contains more protein species (Figure 1F), indicating that the SEC method retains more intrinsic protein species in exosomes during purification. The protein accumulation curve showed that the impurity proteins of MK-exo prepared by the SEC method were ranked lower and no significant difference in positive proteins (Figure 1H), indicating that its heteroprotein content was lower. The proteins linear correlation curve showed that R^2^ was 0.73674, illustrating that the common proteins of MK-exo collected by both methods are highly correlated (Figure 1F). The analysis of the proteome can show that the MK-exo collected by the SEC method is superior. Moreover, the content of whole milk protein of MK-exo prepared by SEC method was significantly lower than that of DC method, and the content of other impurity proteins was low but not obvious (Table 1) This indicates that the content of MK-exo magazine protein prepared by the SEC method is lower, which corroboration the results of proteomics analysis.

**Table 1:** Detection of impurity proteins in MK-exo isolated by DC and SEC.

| Impurity proteins | SEC | DC | P value |
| --- | --- | --- | --- |
| | Mean ± SD | Mean ± SD | |
| Whole milk protein ($\mu\text{g/ml}$) | 3 ± 0.22 | 21.51 ± 14.63 | 0.19 |
| casein ($\mu\text{g/ml}$) | 0.11 ± 0.07 | 2.25 ± 1.53 | 0.07 |
| beta-lactoglobulin ($\mu\text{g/ml}$) | 0 ± 0 | 1.02 ± 0.76 | 0.0801 |
| IgM ($\mu\text{g/ml}$) | 6.29 ± 3.36 | 20.5 ± 28.2 | 0.43 |
| IgG ($\mu\text{g/ml}$) | 0.21 ± 0.09 | 0.53 ± 0.34 | 0.19 |

The above results indicate that we have obtained high-quality MK-exo on a large scale using the SEC method, which can be used for subsequent storage and stability studies.

### 3.2. Detection method validation

To ensure the reliability of our experimental result, the detection method was validated against the *Bioanalytical method validation and study sample analysis M10 published by the International council for harmonisation of technical requirements for harmaceuticals for human use in 2022* to evaluate for linearity and precision et al. [38, 39].

First, Considering the simplicity and onvenience of the assay, NanoFCM was chosen to evaluate the mean and median size of particles (Supplementary table 1-2). Both intra-day and inter-day accuracy were less than 15%, which is qualified (Supplementary table 3-4). And LC-MS/MS was used to detect protein markers and the method proved to be stable (Supplementary Table 5). In addition, in simple terms, the linearity and precision involved in the verification of protein concentration, purity, zeta potential and whole milk protein detection methods are qualified (Supplementary table 6-17). The above results show that the detection method is reliable.

### 3.3. MK-exo stored in PBS at −80℃

Liquid MK-exo can be stored at −80 °C (Munagala et al. 2016, Agrawal et al. 2017). When liquid MK-exo was stored at −80 °C for 8 months, we tested it for morphology, particle size, and purity. During this period, there was no statistically significant difference in morphology and particle size of MK-exo (Figure 2A-B), but its purity had been significantly reduced (Figure 2C). Munagala et al. found no significant change in liquid milk exosome size at −80 storage for 18 months, and our particle size results support this view. But purity result suggests that some physical disruptions that are not visible in electron microscopy and particle size have occurred, and that some proteins exposed have been detected by HPLC. This finding suggests that the physical properties of liquid MK-exo may be unstable after 8 months of storage at −80°C.

**Fig 2.**
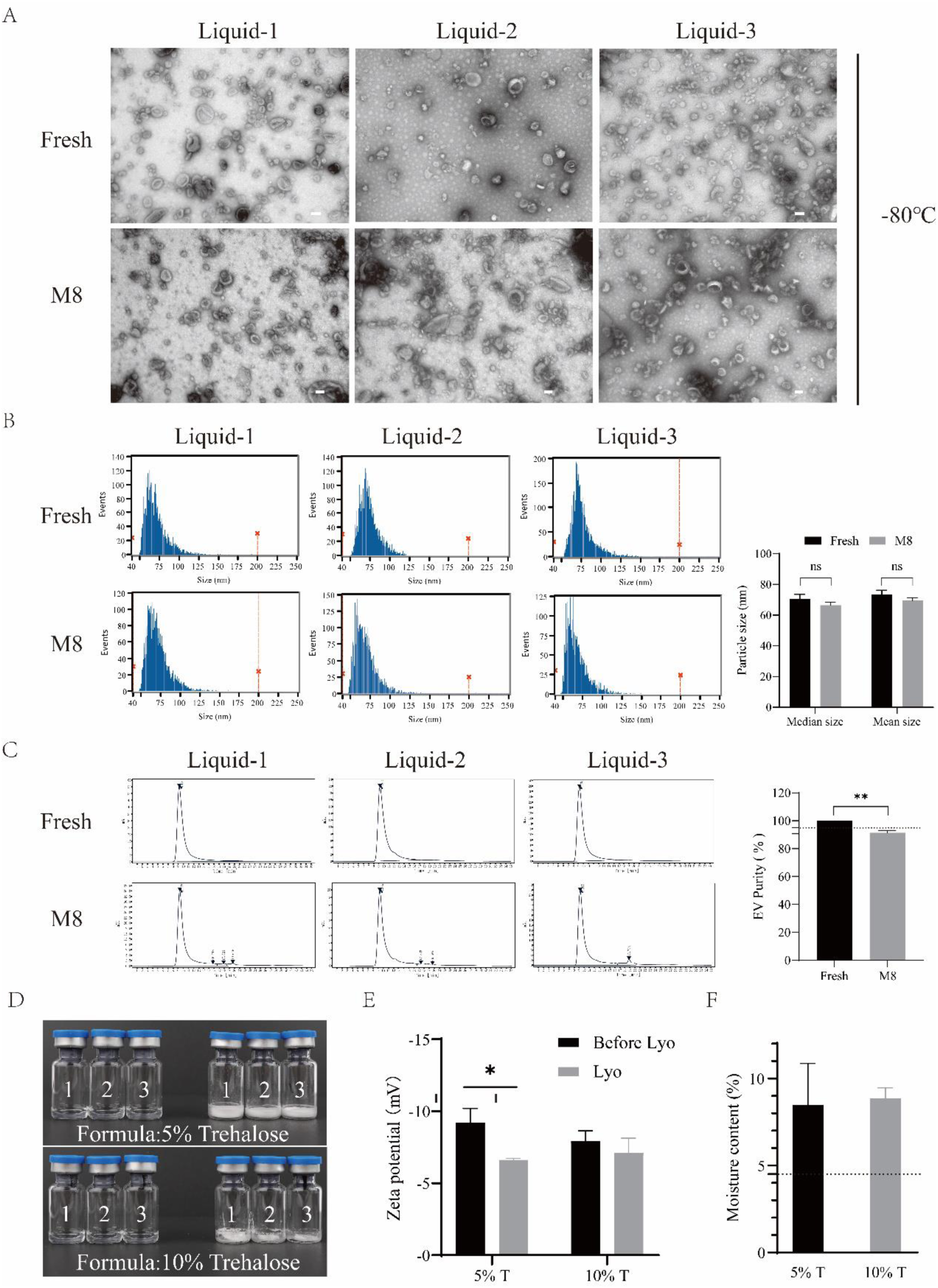
Characterization of liquid MK-exo before and after storage at −80 °C.(A) The main morphological characteristics, particle size analysis and purity analysis of liquid MK-exo; Scale bar = 100 nm; (D) Appearance of lyophilized liquid MK-exo by the process in the literature; (E) Purity analysis of MK-exo before and after lyophilization. The error bars represented the standard deviation of three repetitive experiments. ns: There was no significant difference. \*\**P*<0.01, \**P*<0.05.

### 3.4. Investigation of other lyophilization methods for EVs

To compensate for the shortcomings of −80 °C in terms of long-term storage, MK-exo was lyophilized using the lyophilization methods in the literature(L. et al. 2018, Geeurickx et al. 2019). However, the lyophilized MK-exo is shrinking, collapsed or not cake-like in appearance, and is still sticky (Figure 2D). In the assay we found a statistically significant decrease in zeta potential of MK-exo lyphilized in 5% trehalose, and a slight decrease but no statistical difference in 10% trehalose (Figure 2E). This indicates that MK-exo may have been physically destroyed in lyophilization, exposing some positively charged proteins and causing the potential to rise. And in the moisture test, both prescription samples showed high moisture content (Figure 2F), which indicates that lyophilized is not sufficient and may be risky in later storage.

This suggests that lyophilization methods for other exosomes may not be suitable for MK-exo lyophilization. The lyophilization method for MK-exo needs to be developed.

### 3.5. The lyophilization method of MK-exo is not suitable for MSCs and 293 cell exosomes Lyophilization of MK-exo

In order to obtain qualified lyophilized MK-exo, a lyophilization method suitable for MK-exo was developed and the lyophilized MK-exo was characterized. The 3 batches of MK-exo were lyophilized.

The lyophilized MK-exo has a white cake-like appearance and no shrinkage and collapse occurs. The moisture content of lyophilized MK-exo is 4% (Figure 3A). Meanwhile, MK-exo before and after lyophilization (Lyo) both characterized a classic central depression shape (Figure 3B). The Nano-FCM analysis revealed no difference in particle diameter mean size and median size (Figure 3C). Purity analysis both showed only one peak (Figure 3D), and the zeta potential was no significant difference (Figure 3E). The above indicates that lyophilization has no significant effect on the physical properties of MK-exo.

**Fig 3.**
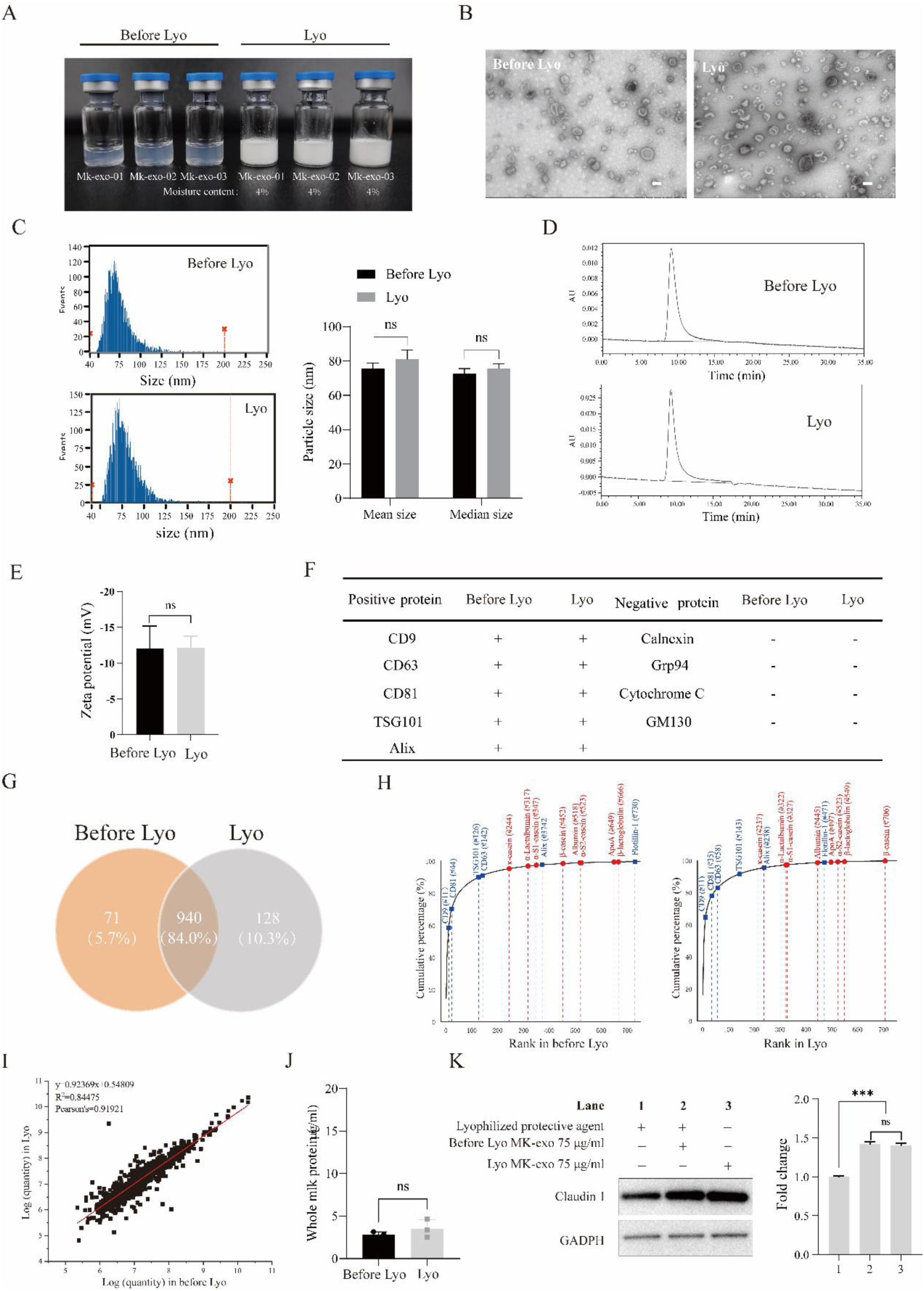
Characterization of liquid and lyophilized MK-exo. (A) The photo of liquid and lyophilizedMK-exo; (B-D) The main morphological features, particle size analysis and proteome Wayne diagram of liquid and lyophilized MK-exo, Scale bar = 100 nm; (E) Condition of milk exosome protein markers of liquid and lyophilized MK-exo.;(F) Protein quantity accumulation analysis between specific surface proteins of exosomes and major milk proteins (impurity protein) of MK-exo before and after lyophilization;(G-I) The protein correlation analysis, purity analysis and zeta potential of liquid and lyophilized MK-exo; (J) The expression of Claudin 1 and GAPDH were determined by western blot and the bands were analyzed based on a grayscale;(K) The content of whole milk protein in liquid and lyophilized MK-exo. The error bars represented the standard deviation of three repetitive experiments. ns: There was no significant difference.\*\*\**P*<0.001.

Furthermore, the marker protein results for MK-exo before and after lyophilization were consistent (Figure 3F). The Venn diagram does not show significant protein differences in MK-exo before and after lyophilization (Figure 1G). And the cumulative curve shows similar protein ordering and heteroprotein content (Figure 3H). The correlation curve also shows a high correlation between the proteins of two (Figure 3I). Meanwhile, there was no significant difference in the protein content of whole milk before and after MK-exo lyophilized (Figure 3J). The results of above showed that there was no significant change in the protein of MK-exo by lyophilization.

Moreover, western blot was used to evaluate the activity of MK-exo before and after lyophilization (Figure 3K). The results of gray band analysis showed that the expression level of Claudin1 protein could be increased by MK-exo before and after lyophilization, and there was no significant difference between them. This showed that lyophilization had no significant effect on the biological activity of MK-exo. The above results showed that MK-exo was successfully lyophilized, which physicochemical properties and biological activity were not destroyed.

Subsequently, the developed MK-exo lyophilization method was used to freeze-dry exosomes of MSCs and 293 cells (Supplementary Fig1A-C), and the purity of the samples after lyophilization decreased significantly (Supplementary Fig1D-G). This further suggests that lyophilization methods for different EVs are not universal. These results suggest that in a range of exosomes, we have developed a lyophilization method that may belong only to MK-exo.

### 3.6. Accelerated experiments of the lyophilized MK-exo

To confirm the stability of MK-exo, ccelerated experiments was designed. After redissolve with PBS, MK-exo were tested at days 0, 14, 30, 60, and 90 of storage (Figure 4A).

**Fig 4.**
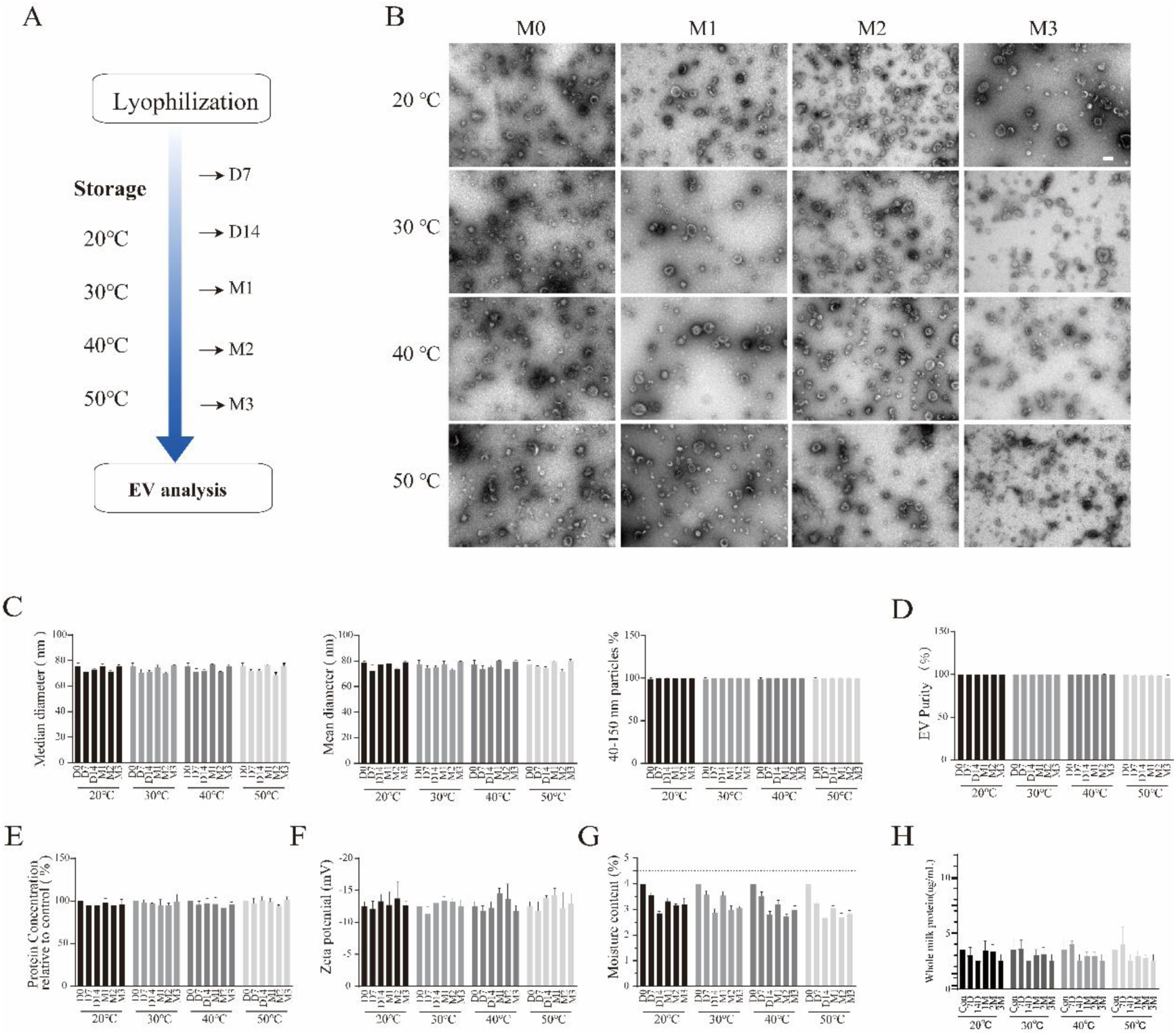
The stability of lyophilized MK-exo in accelerated experiments. (A) Schematic diagram of the accelerated experiment design. (B-H) The analysis of morphology, particle size, purity, protein concentration, zeta potential, moisture content and whole milk protein of lyophilized MK-exo at different temperatures and time points. Scale bar = 500 nm. The error bars represented the standard deviation of three repetitive experiments.

When stored at 50°C for 3 months, MK-exo morphology changed significantly, while other detection sites did not (Figure 4B). There was no significant change in particle size at all detection points, and more than 98% of particles were distributed at 40-150 nm (Figure 4C and Supplementary Fig 2). MK-exo purity stored at 50°C decreased the most, dropping to quality control standards at 3 months (Figure 4D and Supplementary Fig 3). In addition, the results of protein concentration, zeta potential, moisture, and whole milk protein content did not change significantly at each test point and were within quality control standards (Figure 4E-H). Other than that, the marker protein results of MK-exo were all consistent (Supplementary Table 17), and the heat map did not analyze significant differences in the proteome (Supplementary Fig 4).

Moreover, according to the report of Naceur Haouet et al., (Haouet et al. 2018) the shelf life of lyophilized MK-exo at 25 °C was calculated to be 220 days with a concentration reduction of 90% as the qualification criterion. Ich The above results show that lyophilized MK-exo can be stored stable for 3 months at 20, 30 and 40 °C with unchanged physical properties, but will be affected when stored at 50 °C for 3 months. The experiment was done for 3 months, and the actual stable storage time at 20-40 °C may be longer.

### 3.7. Long-term stability experiments of the lyophilized MK-exo

Considering that 2-8 °C is an easily achievable transport and storage temperature, long-term stability experiments for storage at 2-8 °C are further performed (Figure 5A). The experiment is currently in its 15th month.

**Fig 5.**
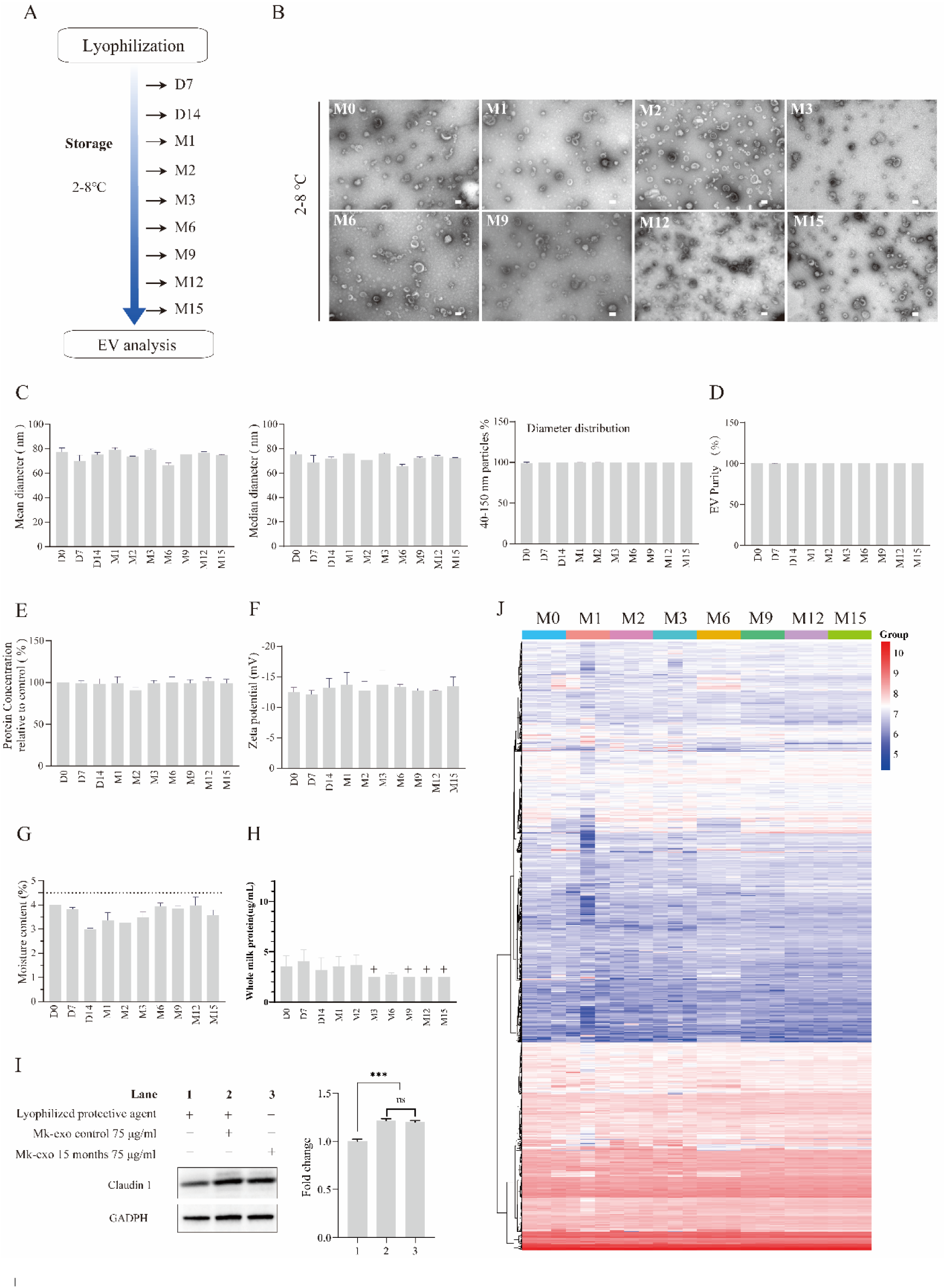
The stability of lyophilized MK-exo in long-term storage at 2-8℃. (A) Schematic diagram of the long-term stability experiment design. (B-H) The analysis of morphology, particle size, purity, protein concentration, zeta potential, moisture content and whole milk protein of lyophilized MK-exo in long-term storage.(I) The expression of Claudin 1 and GAPDH were determined by western blot and the bands were analyzed based on a grayscale;(J) The proteome heatmaps of lyophilized MK-exo in 15 months stored at 2-8℃.

During these 15 months, there were no significant changes in morphology, particle size, purity, protein concentration, zeta potential, moisture and whole milk protein content of MK-exo at all detection point (Figure 5B-H and Supplementary Figure 5-6). Furthermore, the results of western blot showed that the expression level of Claudin1 protein was increased by lyophilized MK-exo before and after 15 months of storage, and there was no significant difference between the two, indicating that the biological activity was stable after storage at 2-8°C for 15 months (Figure 5I). Meanwhile, the marker protein results of MK-exo were consistent across all detection points (Supplementary Table 18), and the heat map did not analyze significant differences in the proteome.

The above results show that the physicochemical properties and biological activity of lyophilized MK-exo do not change significantly after 15months stores at 2-8°C. Thus, lyophilized MK-exo can be stably stored at 2-8 °C for at least 15 months. At the same time, the shelf life at 4 °C was predicted to be 535125936 days using data from accelerated experiments. This prediction results do not exclude factors such as large temperature differences and complex responses.

## 4. DISCUSSION

In this study, we developed a new lyophilization method for MK-exo. Stability experiments have proved that our lyophilized samples can be stored stably for a long time. The accelerated experiments in this paper proved that lyophilized MK-exo can be stored at 20-40 °C for 3 months, which from temperature and duration shows that conventional long-distance transportation is inevitable to achieve. In addition, long-term stability experiments indicating that our lyophilized MK-exo can be stored at 4°C for no less than 15 months. At present, the long-term stability experiment is still going on, and it has been carried out for the 18th month., which gives the lyophilized MK-exo a shelf life that meets expectations.

And we found that liquid MK-exo can indeed be stored at −80°C, but not for too long. Because there was a significant decrease in purity at 8 months, even though no significant change in morphology and particle size of MK-exo. This indicated that the evaluation of exosome rupture may not be limited to morphology and particle size, but also consider purity. Because some changes may not be observed by electron microscopy and particle size, such as the protein passing through the membrane of exosomes to the outside. Freezing at −80°C is the conventional storage method for exosomes (Lorincz et al. 2014, Munagala et al. 2016, Agrawal et al. 2017). However, some reports showed that exosomes are not stable in long-term storage under −80℃. A study by Maroto et al. demonstrated that storage at −80 °C for 4 days changed the morphology of airway exosomes compared to freshly isolated exosomes(Maroto et al. 2017). Analyses of Wu et al. suggested that direct freezing affects the stability of the SKBR3 cells exosome membranes and degrades the samples (Wu et al. 2015). These findings illustrate that −80 °C storage is controversial, possibly because the differences in the source, quality and preparation process of exosomes. Exosomes derived from different cells or biological samples have different components of lipids, RNA, proteins, etc. Different component liposomes lead to different stability of exosome’s membranes. The stability of different RNA and protein to −80 °C may also be different. The potential and proteome results of exosomes from different sources tested in our lab are different, indicating that they do have differences in physical properties and biological activity. Different cell culture methods may also secrete different quality exosomes. Meanwhile, different purification processes result in the harvested samples containing impurities of different compositions and levels. These differences can all lead to different state in exosomes when stored at −80 °C.

Subsequently, we found that the lyophilization method for different EVs is not universal. The lyophilization method of other EVs reported (Geeurickx et al. 2019) were used to freeze-dry MK-exo, but the cake is atrophied, collapsed and viscous with high moisture content and decreased potential. Meanwhile, the lyophilization method of MK-exo we developed was used to freeze-dry MSCs and 293 cells exosomes, and the result was excellent appearance but reduced purity. Among them, the main reasons for the appearance of unqualified may be the lyophilization curve and prescription. Other reasons for poor results may be the differences in the source, quality and preparation process of exosomes as described in the previous paragraph. This difference lead to differences in its biological and physicochemical properties, which is reflected in the lyophilization effect. This leads to challenges in obtaining high-quality and identical exosomes and developing lyophilization methods.

The stable storage of lyophilized MK-exo enables more applications. It can be added to products without heat treatment processes. AndMK-exo are rich in milk fat globule membrane (MFGM) proteins and other nutrients(Reinhardt et al. 2012, Buratta et al. 2023), so adding lyophilized MK-exo to milk powder can help babies develop their brains. Adding it to health supplements can improve the body’s metabolic pattern, strengthen immune function, and improve intestinal flora. And we have investigated that liquid MK-exo can be stored stably at room temperature for 1 month, so MK-exo also expects to be added to milk and yogurt.

