## Supplementary Table&Figure for "Study on the preparation of bovine milk exosomes and the stability of lyophilized powder"

SUPPLEMENTARY TABLE 1 Comparison of results of particle median size detected by TEM and NanoFCM.

SUPPLEMENTARY TABLE 2 Comparison of results of particle mean size detected by TEM and NanoFCM.

SUPPLEMENTARY TABLE 3 The precision of the particle median size of Lyophilized MK-Exo detected by NanoFCM.

SUPPLEMENTARY TABLE 4 The precision of the particle mean size of Lyophilized MK-Exo detected by NanoFCM.

SUPPLEMENTARY TABLE 5 Performance verification for the detection of MK-Exo's surface-tagged protein by LC-MS.

SUPPLEMENTARY TABLE 6 Accuracy verification of protein concentration detection of MK-Exo by kit. (BCA)

SUPPLEMENTARY TABLE 7 Linear verification of protein concentration detection of MK-Exo by kit. (BCA)

SUPPLEMENTARY TABLE 8 Precision verification of protein concentration detection of liquid MK-Exo by kit. (BCA).

SUPPLEMENTARY TABLE 9 Precision verification of protein concentration detection of lyophilized MK-Exo by kit. (BCA)

SUPPLEMENTARY TABLE 10 Precision verification of HPLC detection of liquid MK-Exo.

SUPPLEMENTARY TABLE 11 Linear verification of HPLC detection of liquid MK-Exo.

SUPPLEMENTARY TABLE 12 Precision verification of HPLC detection of lyophilized MK-Exo.

SUPPLEMENTARY TABLE 13 Linear verification of HPLC detection of lyophilized MK-Exo.

SUPPLEMENTARY TABLE 14 Precision verification of zeta potential detection of liquid MK-Exo.

SUPPLEMENTARY TABLE 15 Precision verification of zeta potential detection of lyophilized MK-Exo.

SUPPLEMENTARY TABLE 16 Precision verification of whole milk protein detection of lyophilized MK-Exo by kit.

SUPPLEMENTARY TABLE 17 Linear verification of whole milk protein detection of lyophilized MK-Exo.

SUPPLEMENTARY TABLE18 MK-Exo protein marker test for accelerate experimentation.

SUPPLEMENTARY FIGURE 1 MSCs and 293 cell exosomes were identified and lyophilized.

SUPPLEMENTARY FIGURE 2 Particle size distribution of lyophilized MK-Exo after redissolve in accelerated experiments was analyzed.

SUPPLEMENTARY FIGURE 3 The purity of lyophilized MK-Exo after redissolve in accelerated experiments was detected.

SUPPLEMENTARY FIGURE 4 Heat map of the proteome of lyophilized MK-Exo in accelerated experiments was made.

SUPPLEMENTARY FIGURE 5 Particle size distribution of lyophilized MK-Exo after redissolve in long-term stabilization experiments was detected.

SUPPLEMENTARY FIGURE 6 Purity of lyophilized MK-Exo after redissolve in long-term stabilization experiments was detected.

SUPPLEMENTARY TABLE 1 Comparison of results of particle median size detected by TEM and NanoFCM.

| Time | TEM | | NanoFCM | | P Value |
| --- | --- | --- | --- | --- | --- |
|  | Median size (nm) | average | Median size (nm) | average |  |
| D1 | 75.76 | 73.18 | 73.25 | 73.75 | 0.035875 |
|  | 68.96 |  | 73.75 |  |  |
|  | 74.83 |  | 74.25 |  |  |
| D2 | 69.62 | 70.32 | 76.25 | 75.75 |  |
|  | 70.78 |  | 75.75 |  |  |
|  | 70.56 |  | 75.25 |  |  |
| D3 | 70.25 | 69.36 | 78.75 | 76.42 |  |
|  | 67.71 |  | 75.75 |  |  |
|  | 70.12 |  | 74.75 |  |  |

SUPPLEMENTARY TABLE 2 Comparison of results of particle mean size detected by TEM and NanoFCM.

| Time | TEM | | NanoFCM | | P Value |
| --- | --- | --- | --- | --- | --- |
|  | Mean size | average | Mean size | average |  |
|  | (nm) |  | (nm) |  |  |
| D1 | 81.44 | 80.08 | 76.12 | 76.61 | 0.3816 |
|  | 75.52 |  | 76.60 |  |  |
|  | 83.28 |  | 77.11 |  |  |
| D2 | 74.36 | 76.53 | 79.33 | 78.83 |  |
|  | 77.43 |  | 78.66 |  |  |
|  | 77.81 |  | 78.50 |  |  |
| D3 | 76.65 | 76.06 | 82.48 | 80.43 |  |
|  | 74.71 |  | 79.70 |  |  |
|  | 76.82 |  | 79.12 |  |  |

SUPPLEMENTARY TABLE 3 The precision of the particle median size of Lyophilized MK-Exo detected by NanoFCM.

| Time | Repetition | Median size（nm) | SD | Average(nm) | Intra-day  Precision(%) | Inter-day  Precision(%) |
| --- | --- | --- | --- | --- | --- | --- |
| D1 | 1 | 77.25 | 0.52 | 77.58 | 0.67 | 4.21 |
|  | 2 | 78.25 |  |  |  |  |
|  | 3 | 77.25 |  |  |  |  |
|  | 4 | 78.25 |  |  |  |  |
|  | 5 | 77.25 |  |  |  |  |
|  | 6 | 77.25 |  |  |  |  |
| D2 | 1 | 71.25 | 0.71 | 71.25 | 0.99 |  |
|  | 2 | 70.75 |  |  |  |  |
|  | 3 | 72.25 |  |  |  |  |
|  | 4 | 71.75 |  |  |  |  |
|  | 5 | 71.25 |  |  |  |  |
|  | 6 | 70.25 |  |  |  |  |
| D3 | 1 | 70.25 | 0.94 | 71.50 | 1.31 |  |
|  | 2 | 72.75 |  |  |  |  |
|  | 3 | 71.25 |  |  |  |  |
|  | 4 | 71.75 |  |  |  |  |
|  | 5 | 72.25 |  |  |  |  |
|  | 6 | 70.75 |  |  |  |  |

SUPPLEMENTARY TABLE 4 The precision of the particle mean size of Lyophilized MK-Exo detected by NanoFCM.

| Time | Repetition | Mean size（nm) | SD | Average(nm) | Intra-day Precision(%) | Inter-day Precision(%) |
| --- | --- | --- | --- | --- | --- | --- |
| D1 | 1 | 81.11 | 0.71 | 81.52 | 0.87 | 4.43 |
|  | 2 | 82.55 |  |  |  |  |
|  | 3 | 81.43 |  |  |  |  |
|  | 4 | 82.24 |  |  |  |  |
|  | 5 | 80.81 |  |  |  |  |
|  | 6 | 80.99 |  |  |  |  |
| D2 | 1 | 74.30 | 0.78 | 74.63 | 1.04 |  |
|  | 2 | 74.35 |  |  |  |  |
|  | 3 | 75.63 |  |  |  |  |
|  | 4 | 75.56 |  |  |  |  |
|  | 5 | 74.20 |  |  |  |  |
|  | 6 | 73.75 |  |  |  |  |
| D3 | 1 | 73.31 | 1.09 | 74.77 | 1.46 |  |
|  | 2 | 76.20 |  |  |  |  |
|  | 3 | 74.44 |  |  |  |  |
|  | 4 | 75.19 |  |  |  |  |
|  | 5 | 75.62 |  |  |  |  |
|  | 6 | 73.87 |  |  |  |  |

SUPPLEMENTARY TABLE 5 Performance verification for the detection of MK-Exo's surface-tagged protein by LC-MS.

|  | | MK-Exo-N1 | MK-Exo-N2 | MK-Exo-N3 | MK-Exo-N4 | MK-Exo-N5 | MK-Exo-N6 | MK-Exo-N7 | Blank 1 | Blank 2 |
| --- | --- | --- | --- | --- | --- | --- | --- | --- | --- | --- |
| Exosome marker proteins | CD9(P30932) | + | + | + | + | + | + | + | - | - |
|  | CD63(Q9XSK2) | + | + | + | + | + | + | + | - | - |
|  | CD81(Q3ZCD0) | + | + | + | + | + | + | + | - | - |
|  | TSG101(Q99816) | + | + | + | + | + | + | + | - | - |
|  | Alix(E1BKM4) | + | + | + | + | + | + | + | - | - |
|  | Flotillin1(Q08DN8) | + | - | + | + | + | - | + | - | - |
|  | Flotillin2(A6QLR4) | + | + | + | - | - | - | + | - | - |
|  | AQP2(P79099) | - | - | - | - | - | - | - | - | - |
|  | Lamp2(P13473) | - | - | - | - | - | - | - | - | - |
| Exosome-negative proteins | Calnexin((P27824) | - | - | - | - | - | - | - | - | - |
|  | Grp94(Q95M18) | - | - | - | - | - | - | - | - | - |
|  | Cytochrome C（P00125） | - | - | - | - | - | - | - | - | - |
|  | GM130 | - | - | - | - | - | - | - | - | - |

SUPPLEMENTARY TABLE 6 Accuracy verification of protein concentration detection of MK-Exo by kit. (BCA)

| Repetition1  (ug/mL) | Repetition2  (ug/mL) | mean value(ug/mL) | spiked value(ug/mL) | Calculated value(ug/mL) | Measured value(ug/mL) | recovery % | RSD(%) |
| --- | --- | --- | --- | --- | --- | --- | --- |
| 112.845 | 104.045 | 108.445 | 217.669 | 163.057 | 150.271 | 92.16 | 2.79 |
| 95.815 | 99.175 | 97.495 | 217.669 | 157.582 | 154.6975 | 98.17 |  |
| 105.754 | 114.088 | 109.921 | 217.669 | 163.795 | 159.589 | 97.43 |  |
| 109.895 | 108.652 | 109.2735 | 217.669 | 163.47125 | 157.57 | 96.39 |  |
| 115.33 | 113.01 | 114.17 | 217.669 | 165.9195 | 162.3585 | 97.85 |  |
| 115.589 | 118.074 | 116.8315 | 217.669 | 167.25025 | 155.008 | 92.68 |  |

SUPPLEMENTARY TABLE 7 Linear verification of protein concentration detection of MK-Exo by kit. (BCA)

| Time | Linear range (ug/mL) | R^2^ | Calibration Curve Equation |
| --- | --- | --- | --- |
| D1 | 23.125-370 | 0.9981 | y = 1.0013x - 1.995 |
| D2 |  | 0.9998 | y = 1.0153x - 2.1313 |
| D3 |  | 0.9985 | y = 1.0584x - 7.7507 |

SUPPLEMENTARY TABLE 8 Precision verification of protein concentration detection of liquid MK-Exo by kit. (BCA).

| Time | Repetition | Calculated Conc.(μg/ml) | Average (μg/ml) | SD | Intra-day Precision% | Inter-day Precision% |
| --- | --- | --- | --- | --- | --- | --- |
| D1 | 1 | 251.965 | 253.35 | 3.96 | 1.56 | 2.14 |
|  | 2 | 249.659 |  |  |  |  |
|  | 3 | 249.429 |  |  |  |  |
|  | 4 | 252.657 |  |  |  |  |
|  | 5 | 257.960 |  |  |  |  |
|  | 6 | 258.421 |  |  |  |  |
| D2 | 1 | 243.477 | 247.29 | 3.96 | 1.60 |  |
|  | 2 | 245.609 |  |  |  |  |
|  | 3 | 242.648 |  |  |  |  |
|  | 4 | 249.398 |  |  |  |  |
|  | 5 | 250.050 |  |  |  |  |
|  | 6 | 252.537 |  |  |  |  |
| D3 | 1 | 240.378 | 245.33 | 4.81 | 1.96 |  |
|  | 2 | 241.572 |  |  |  |  |
|  | 3 | 242.012 |  |  |  |  |
|  | 4 | 246.222 |  |  |  |  |
|  | 5 | 249.678 |  |  |  |  |
|  | 6 | 252.129 |  |  |  |  |

SUPPLEMENTARY TABLE 9 Precision verification of protein concentration detection of lyophilized MK-Exo by kit. (BCA)

| Time | Repetition | Calculated Conc.(μg/ml) | Average (μg/ml) | SD | Intra-day  Precision% | Inter-day  Precision% |
| --- | --- | --- | --- | --- | --- | --- |
| D1 | 1 | 276.511 | 281.71 | 4.73 | 1.68 | 4.39 |
|  | 2 | 279.093 |  |  |  |  |
|  | 3 | 279.978 |  |  |  |  |
|  | 4 | 281.527 |  |  |  |  |
|  | 5 | 282.855 |  |  |  |  |
|  | 6 | 290.305 |  |  |  |  |
| D2 | 1 | 283.359 | 300.10 | 10.82 | 3.61 |  |
|  | 2 | 314.885 |  |  |  |  |
|  | 3 | 308.565 |  |  |  |  |
|  | 4 | 298.510 |  |  |  |  |
|  | 5 | 298.056 |  |  |  |  |
|  | 6 | 297.247 |  |  |  |  |
| Da3 | 1 | 271.388 | 288.19 | 14.26 | 4.95 |  |
|  | 2 | 278.919 |  |  |  |  |
|  | 3 | 309.119 |  |  |  |  |
|  | 4 | 297.22 |  |  |  |  |
|  | 5 | 294.056 |  |  |  |  |
|  | 6 | 278.467 |  |  |  |  |

SUPPLEMENTARY TABLE 10 Precision verification of HPLC detection of liquid MK-Exo.

| Time | Repetition | Calculated Aera | Average | SD | Intra-day Precision (%) | Inter-day Precision (%) |
| --- | --- | --- | --- | --- | --- | --- |
| D1 | 1 | 968.4896 | 1061.5015 | 52.50 | 4.95 | 5.55 |
|  | 2 | 1045.726 |  |  |  |  |
|  | 3 | 1095.684 |  |  |  |  |
|  | 4 | 1092.462 |  |  |  |  |
|  | 5 | 1113.347 |  |  |  |  |
|  | 6 | 1053.3006 |  |  |  |  |
| D2 | 1 | 998.104 | 1086.8423 | 50.21 | 4.62 |  |
|  | 2 | 1059.590 |  |  |  |  |
|  | 3 | 1095.040 |  |  |  |  |
|  | 4 | 1127.969 |  |  |  |  |
|  | 5 | 1118.260 |  |  |  |  |
|  | 6 | 1122.089 |  |  |  |  |
| D3 | 1 | 921.379 | 1010.2237 | 51.85 | 5.13 |  |
|  | 2 | 981.444 |  |  |  |  |
|  | 3 | 1012.190 |  |  |  |  |
|  | 4 | 1035.936 |  |  |  |  |
|  | 5 | 1053.036 |  |  |  |  |
|  | 6 | 1057.3571 |  |  |  |  |

SUPPLEMENTARY TABLE 11 Linear verification of HPLC detection of liquid MK-Exo.

| Time | Linear range(particles/mL) | R^2^ | Calibration Curve Equation |
| --- | --- | --- | --- |
| D1 | 6.39E+6-6.39E+7 | 0.9963 | y = 1.0223x - 74.238 |
| D2 | 2.84E+6-6.06E+7 | 0.9972 | y = 1.0165x - 66.779 |
| D3 | 7.54E+5-1.51E+8 | 0.9984 | y = 1.0173x - 52.864 |

SUPPLEMENTARY TABLE 12 Precision verification of HPLC detection of lyophilized MK-Exo.

| Time | Repetition | Calculated Aera | Average | SD | Intra-day  Precision (%) | Inter-day  Precision (%) |
| --- | --- | --- | --- | --- | --- | --- |
| D1 | 1 | 1600.45 | 1841.74 | 155.16 | 8.42 | 8.56 |
|  | 2 | 1735.17 |  |  |  |  |
|  | 3 | 1836.26 |  |  |  |  |
|  | 4 | 1886.95 |  |  |  |  |
|  | 5 | 1970.77 |  |  |  |  |
|  | 6 | 2020.82 |  |  |  |  |
| D2 | 1 | 1781.51 | 1977.58 | 133.28 | 6.74 |  |
|  | 2 | 1892.01 |  |  |  |  |
|  | 3 | 1941.33 |  |  |  |  |
|  | 4 | 2013.81 |  |  |  |  |
|  | 5 | 2099.04 |  |  |  |  |
|  | 6 | 2137.81 |  |  |  |  |
| D3 | 1 | 1567.74 | 1816.46 | 167.31 | 9.21 |  |
|  | 2 | 1665.67 |  |  |  |  |
|  | 3 | 1818.28 |  |  |  |  |
|  | 4 | 1914.95 |  |  |  |  |
|  | 5 | 1944.78 |  |  |  |  |
|  | 6 | 1987.33 |  |  |  |  |

SUPPLEMENTARY TABLE 13 Linear verification of HPLC detection of lyophilized MK-Exo.

| Time | Linear range(particles/mL) | R^2^ | Calibration Curve Equation |
| --- | --- | --- | --- |
| D 1 | 1.39E+7-4.45E+8 | 0.9982 | y = 1.0118x - 106.83 |
| D 2 | 1.47E+7-4.70E+8 | 0.9989 | y = 1.0281x - 159.49 |
| D 3 | 1.42E+7-4.55E+8 | 0.9971 | y = 1.0259x - 176.24 |

SUPPLEMENTARY TABLE 14 Precision verification of zeta potential detection of liquid MK-Exo.

| Time | Repetition | Zeta potential（mV） | Average（mV） | SD | Intra-day  Precision (%) | Inter-day  Precision (%) |
| --- | --- | --- | --- | --- | --- | --- |
| D1 | 1 | -17.54 | -15.90 | 1.026 | 6.450 | 9.143 |
|  | 2 | -15.58 |  |  |  |  |
|  | 3 | -16.54 |  |  |  |  |
|  | 4 | -15.91 |  |  |  |  |
|  | 5 | -14.69 |  |  |  |  |
|  | 6 | -15.13 |  |  |  |  |
| D2 | 1 | -12.39 | -13.85 | 1.049 | 7.574 |  |
|  | 2 | -15.58 |  |  |  |  |
|  | 3 | -14.04 |  |  |  |  |
|  | 4 | -14.09 |  |  |  |  |
|  | 5 | -13.36 |  |  |  |  |
|  | 6 | -13.64 |  |  |  |  |
| D3 | 1 | -17.47 | -15.66 | 1.155 | 7.375 |  |
|  | 2 | -14.44 |  |  |  |  |
|  | 3 | -16.55 |  |  |  |  |
|  | 4 | -15.53 |  |  |  |  |
|  | 5 | -15.32 |  |  |  |  |
|  | 6 | -14.67 |  |  |  |  |

SUPPLEMENTARY TABLE 15 Precision verification of zeta potential detection of lyophilized MK-Exo.

| Time | Repetition | Zeta potential（mV） | Average（mV） | SD | Intra-day  Precision (%) | Inter-day  Precision (%) |
| --- | --- | --- | --- | --- | --- | --- |
| D1 | 1 | -12.57 | -12.50 | 0.49 | 3.93 | 5.25 |
|  | 2 | -12.50 |  |  |  |  |
|  | 3 | -13.15 |  |  |  |  |
|  | 4 | -11.81 |  |  |  |  |
|  | 5 | -12.10 |  |  |  |  |
|  | 6 | -12.88 |  |  |  |  |
| D2 | 1 | -13.55 | -12.94 | 0.57 | 4.44 |  |
|  | 2 | -12.99 |  |  |  |  |
|  | 3 | -13.19 |  |  |  |  |
|  | 4 | -12.07 |  |  |  |  |
|  | 5 | -13.38 |  |  |  |  |
|  | 6 | -12.43 |  |  |  |  |
| D3 | 1 | -13.99 | -12.67 | 0.91 | 7.18 |  |
|  | 2 | -12.05 |  |  |  |  |
|  | 3 | -11.60 |  |  |  |  |
|  | 4 | -12.11 |  |  |  |  |
|  | 5 | -12.90 |  |  |  |  |
|  | 6 | -13.38 |  |  |  |  |

SUPPLEMENTARY TABLE 16 Precision verification of whole milk protein detection of lyophilized MK-Exo by kit.

| Time | Repetition | Measured value(ug/ml) | Average  (ug/ml) | SD | Intra-day  Precision (%) | Inter-day  Precision (%) |
| --- | --- | --- | --- | --- | --- | --- |
| D1 | 1 | 4.77 | 4.80 | 0.12 | 2.54 | 4.52 |
|  | 2 | 4.98 |  |  |  |  |
|  | 3 | 4.77 |  |  |  |  |
|  | 4 | 4.88 |  |  |  |  |
|  | 5 | 4.64 |  |  |  |  |
|  | 6 | 4.71 |  |  |  |  |
| D2 | 1 | 4.94 | 4.71 | 0.24 | 5.13 |  |
|  | 2 | 4.5 |  |  |  |  |
|  | 3 | 4.94 |  |  |  |  |
|  | 4 | 4.51 |  |  |  |  |
|  | 5 | 4.91 |  |  |  |  |
|  | 6 | 4.46 |  |  |  |  |
| D3 | 1 | 4.43 | 4.51 | 0.17 | 3.67 |  |
|  | 2 | 4.38 |  |  |  |  |
|  | 3 | 4.81 |  |  |  |  |
|  | 4 | 4.54 |  |  |  |  |
|  | 5 | 4.53 |  |  |  |  |
|  | 6 | 4.36 |  |  |  |  |

SUPPLEMENTARY TABLE 17 Linear verification of whole milk protein detection of lyophilized MK-Exo.

| Time | Linear range(mg/kg) | R^2^ | Calibration Curve Equation |
| --- | --- | --- | --- |
| D 1 | 0-67.5 | 0.99975 | y=(4.23074-0.13722)/[1+(x/37.31774)^(-1.00135)]+0.13722 |
| D 2 |  | 0.99983 | y=(4.25922-0.13268)/[1+(x/42.61862)^(-0.99218)]+0.13268 |
| D 3 |  | 0.99990 | y=(4.02925-0.13592)/[1+(x/38.74027)^(-1.03087)]+0.13592 |

SUPPLEMENTARY TABLE18 MK-Exo protein marker test for accelerate experimentation.

| **Lot** | | **Mk-Exo 01** | | | | **Mk-Exo 02** | | | | **Mk-Exo 03** | | | |
| --- | --- | --- | --- | --- | --- | --- | --- | --- | --- | --- | --- | --- | --- |
| Temperature ℃ | | 20 | 30 | 40 | 50 | 20 | 30 | 40 | 50 | 20 | 30 | 40 | 50 |
| Positive protein | CD9 | **+** | **+** | **+** | **+** | **+** | **+** | **+** | **+** | **+** | **+** | **+** | **+** |
|  | CD63 | **+** | **+** | **+** | **+** | **+** | **+** | **+** | **+** | **+** | **+** | **+** | **+** |
|  | CD81 | **+** | **+** | **+** | **+** | **+** | **+** | **+** | **+** | **+** | **+** | **+** | **+** |
|  | TSG101 | **+** | **+** | **+** | **+** | **+** | **+** | **+** | **+** | **+** | **+** | **+** | **+** |
|  | Lamp2 | **+** | **+** | **+** | **+** | **+** | **+** | **+** | **+** | **+** | **+** | **+** | **+** |
| Negative protein | Calnexin | **-** | **-** | **-** | **-** | **-** | **-** | **-** | **-** | **-** | **-** | **-** | **-** |
|  | Grp94 | **-** | **-** | **-** | **-** | **-** | **-** | **-** | **-** | **-** | **-** | **-** | **-** |
|  | Cytochrome C | **-** | **-** | **-** | **-** | **-** | **-** | **-** | **-** | **-** | **-** | **-** | **-** |
|  | GM130 | **-** | **-** | **-** | **-** | **-** | **-** | **-** | **-** | **-** | **-** | **-** | **-** |

SUPPLEMENTARY TABLE 19 MK-Exo protein marker test for long-term storage experimentation.

| **Lot** | | **Mk-Exo-01** | | | | | | | **Mk-Exo-02** | | | | | | | **Mk-Exo-03** | | | | | | |
| --- | --- | --- | --- | --- | --- | --- | --- | --- | --- | --- | --- | --- | --- | --- | --- | --- | --- | --- | --- | --- | --- | --- |
| Time ( Month ) | | 1 | 2 | 3 | 6 | 9 | 12 | 15 | 1 | 2 | 3 | 6 | 9 | 12 | 15 | 1 | 2 | 3 | 6 | 9 | 12 | 15 |
| Positive protein | CD9 | **+** | **+** | **+** | **+** | **+** | **+** | **+** | **+** | **+** | **+** | **+** | **+** | **+** | **+** | **+** | **+** | **+** | **+** | **+** | **+** | **+** |
|  | CD63 | **+** | **+** | **+** | **+** | **+** | **+** | **+** | **+** | **+** | **+** | **+** | **+** | **+** | **+** | **+** | **+** | **+** | **+** | **+** | **+** | **+** |
|  | CD81 | **+** | **+** | **+** | **+** | **+** | **+** | **+** | **+** | **+** | **+** | **+** | **+** | **+** | **+** | **+** | **+** | **+** | **+** | **+** | **+** | **+** |
|  | TSG101 | **+** | **+** | **+** | **+** | **+** | **+** | **+** | **+** | **+** | **+** | **+** | **+** | **+** | **+** | **+** | **+** | **+** | **+** | **+** | **+** | **+** |
|  | Lamp2 | **+** | **+** | **+** | **+** | **+** | **+** | **+** | **+** | **+** | **+** | **+** | **+** | **+** | **+** | **+** | **+** | **+** | **+** | **+** | **+** | **+** |
| Negative protein | Calnexin | **-** | **-** | **-** | **-** | **-** | **-** | **-** | **-** | **-** | **-** | **-** | **-** | **-** | **-** | **-** | **-** | **-** | **-** | **-** | **-** | **-** |
|  | Grp94 | **-** | **-** | **-** | **-** | **-** | **-** | **-** | **-** | **-** | **-** | **-** | **-** | **-** | **-** | **-** | **-** | **-** | **-** | **-** | **-** | **-** |
|  | Cytochrome C | **-** | **-** | **-** | **-** | **-** | **-** | **-** | **-** | **-** | **-** | **-** | **-** | **-** | **-** | **-** | **-** | **-** | **-** | **-** | **-** | **-** |
|  | GM130 | **-** | **-** | **-** | **-** | **-** | **-** | **-** | **-** | **-** | **-** | **-** | **-** | **-** | **-** | **-** | **-** | **-** | **-** | **-** | **-** | **-** |

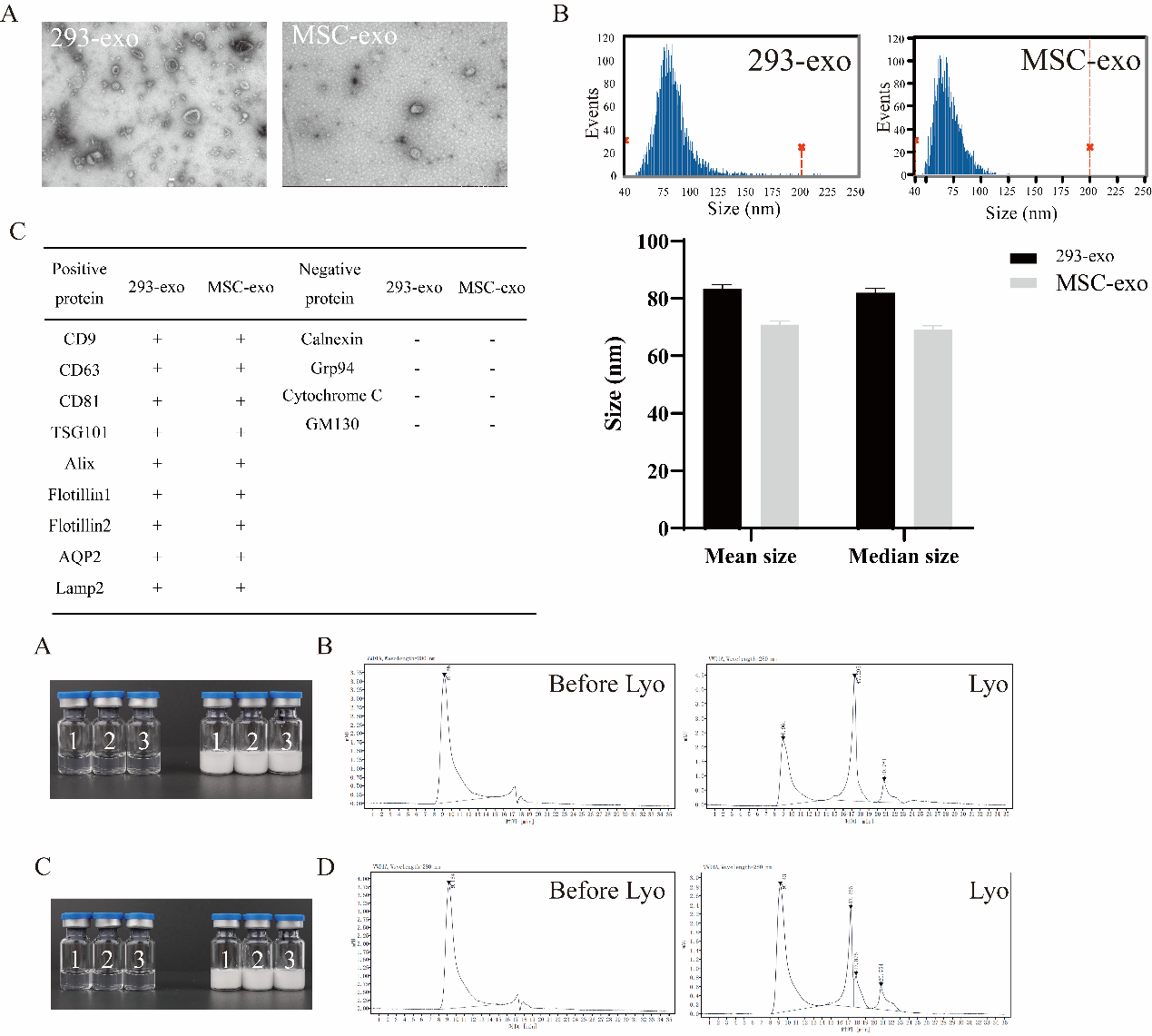

SUPPLEMENTARY FIGURE 1 MSCs and 293 cell exosomes were identified and lyophilized. (A) Exosomes of 293 cells were lyophilized using MK-Exo's lyophilized method and photographed; (B) Purity of 293 cells exosomes before and after lyophilization were analyzed; (C) Exosomes of MSC were lyophilized using MK-Exo's lyophilized method and photographed; (D) Purity of MSC exosomes before and after lyophilization were analyzed.

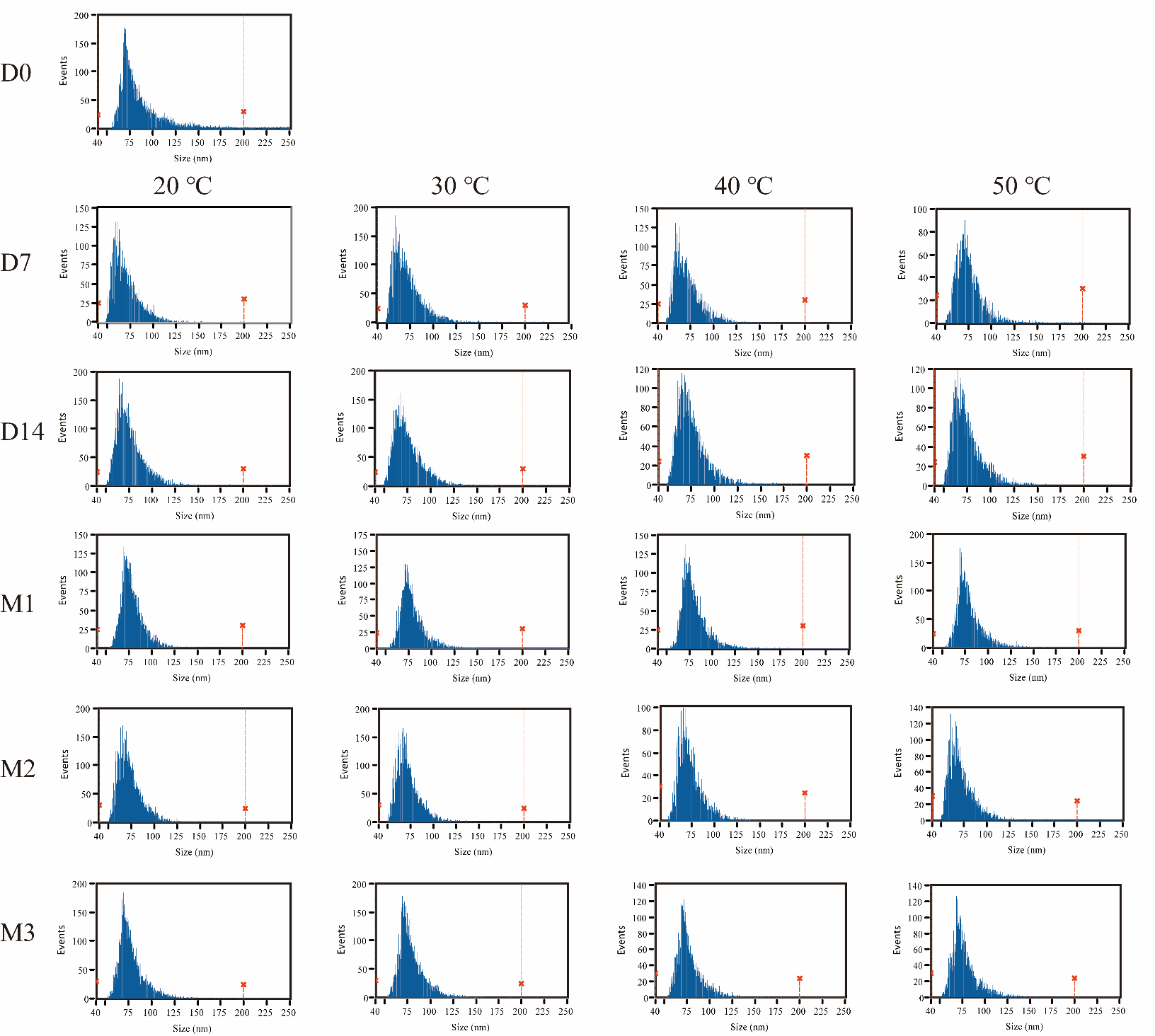

SUPPLEMENTARY FIGURE 2 Particle size distribution of lyophilized MK-Exo after redissolve in accelerated experiments was analyzed.

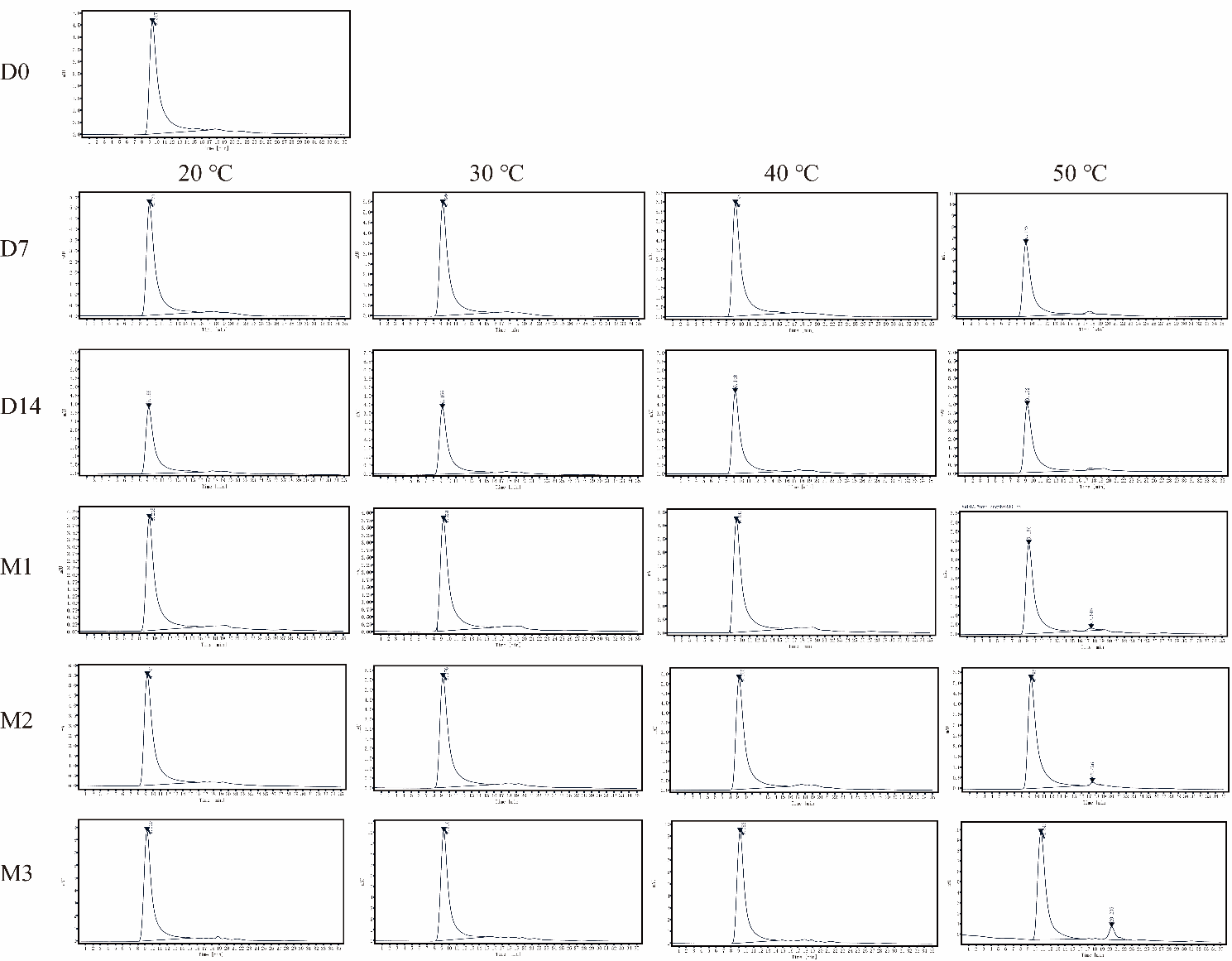

SUPPLEMENTARY FIGURE 3 The purity of lyophilized MK-Exo after redissolve in accelerated experiments was detected.

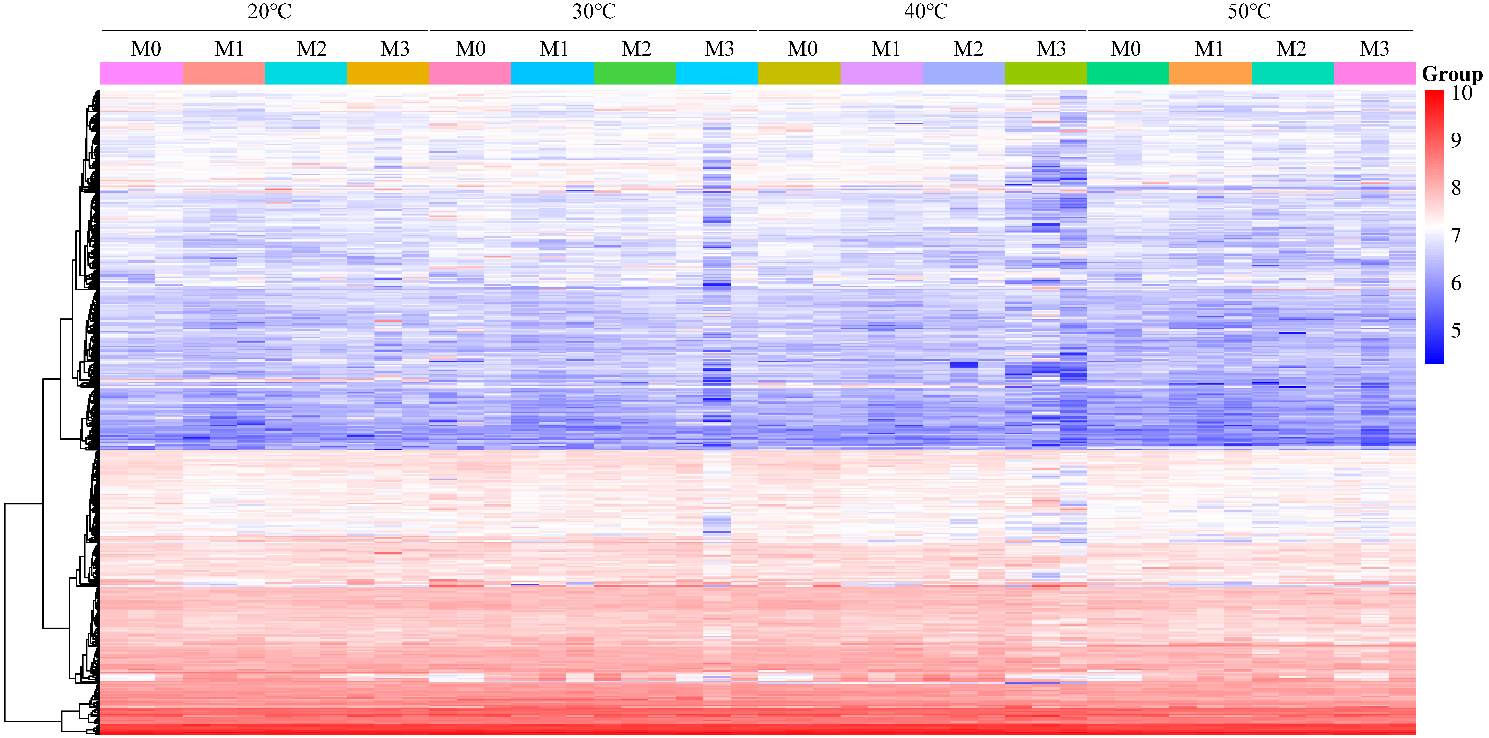

SUPPLEMENTARY FIGURE 4 Heat map of the proteome of lyophilized MK-Exo in accelerated experiments was made.

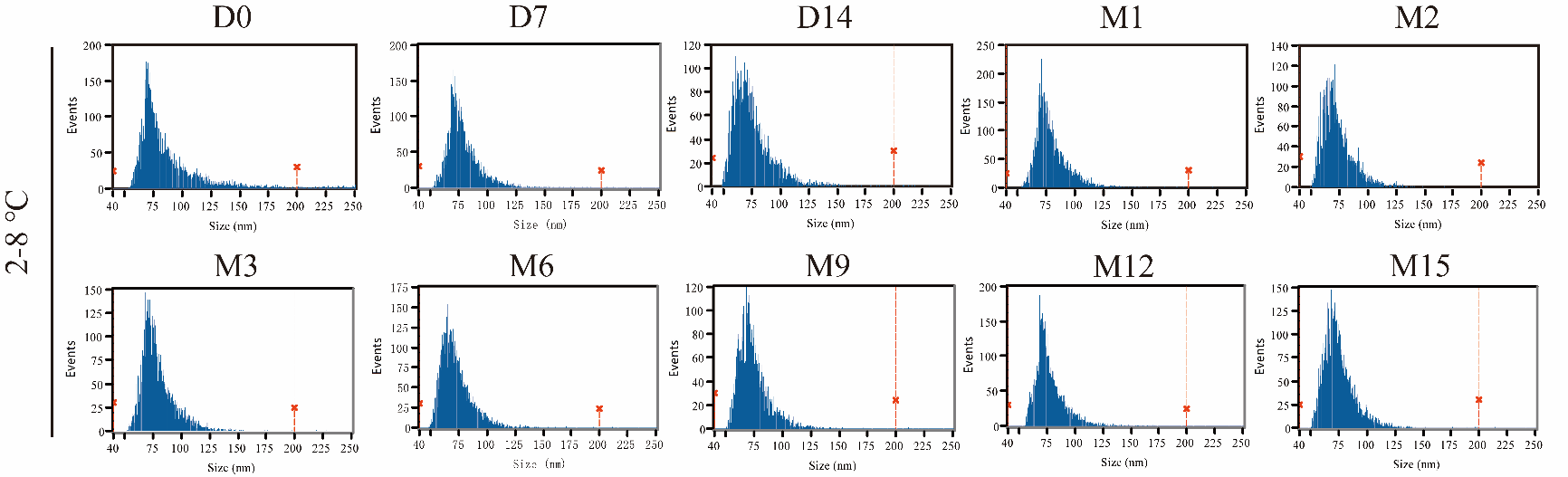

SUPPLEMENTARY FIGURE 5 Particle size distribution of lyophilized MK-Exo after redissolve in long-term stabilization experiments was detected.

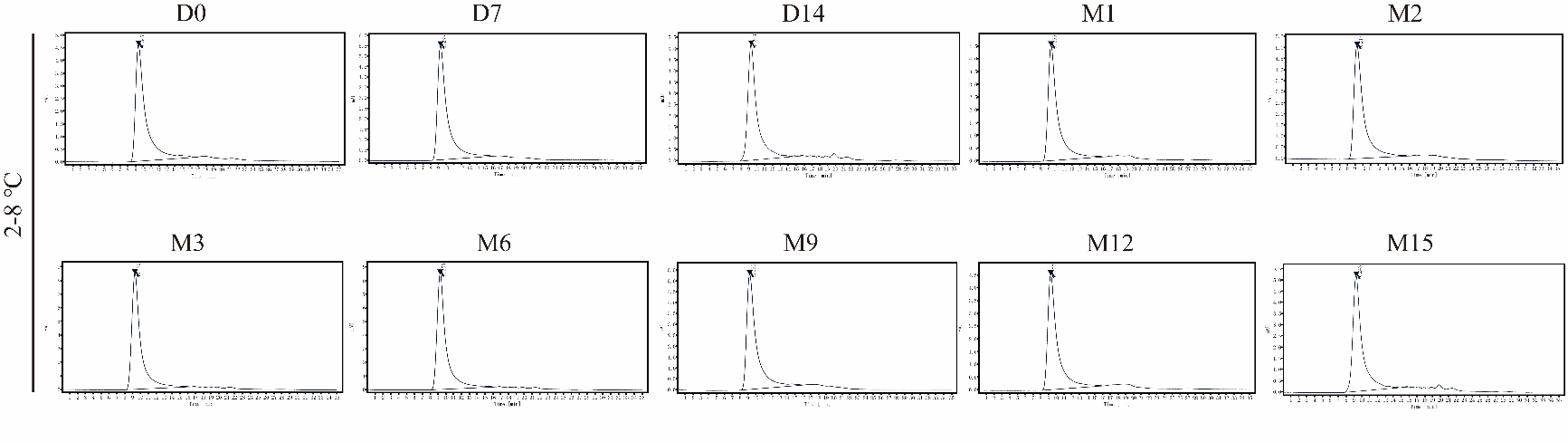

SUPPLEMENTARY FIGURE 6 Purity of lyophilized MK-Exo after redissolve in long-term stabilization experiments was detected.
